# Defective engram allocation contributes to impaired fear memory performance in Down syndrome

**DOI:** 10.1101/2023.01.11.523460

**Authors:** Álvaro Fernández-Blanco, Alfonsa Zamora-Moratalla, Miguel Sabariego-Navarro, Mara Dierssen

## Abstract

Down syndrome (DS) is the most common genetic form of intellectual disability (ID). The cellular and molecular mechanisms contributing to ID in DS are not completely understood. Recent evidence indicates that a given memory is encoded by sparsely distributed neurons, highly activated during learning, the engram cells. Intriguingly, mechanisms that are of paramount importance for engram formation are impaired in DS. Here we explored engram formation in a DS mouse model, the Ts65Dn and we found a reduced number of engram cells in the dentate gyrus (DG), suggesting reduced neuronal allocation to engrams. We also show that trisomic engram cells present reduced number of mature spines than WT engram cells and their excitability is not enhanced during memory recall. In fact, activation of engram cells using a chemogenetic approach does not recover memory deficits in Ts65Dn. Altogether, our findings suggest that perturbations in engram neurons may play a significant role in memory alterations in DS.

## Introduction

Down syndrome (DS) is the most common genetic cause of intellectual disability and results from the presence of an extra copy or major portion of HSA21. Hippocampal-dependent memory functions such as short-term and working memory are particularly affected in individuals with DS [1,2]. Even though research on animal models of DS have revealed extensive cellular and molecular mechanisms that might contribute to DS memory deficits, studies have relied on basic neural functions, such as synaptic transmission, biochemical aspects, ion channel malfunction, or synaptic plasticity [3].

Recently, new molecular and genetic methods such as activity-dependent tagging along with chemo- and optogenetics have provided unprecedented opportunities to visualize and manipulate the neural correlates of memory [4,5]. With these new techniques, different research groups have demonstrated that a sparse assembly of neurons that activate at the time of contextual fear-conditioning (CFC) training are sufficient and necessary for subsequent memory retrieval [5–7]. These neurons, called engram cells, are thought to be the neural correlates of memory. Engram cells activate during learning and undergo molecular and structural changes that enable them to reactivate by recall cues [7,8]. Engrams have been extensively studied in the hippocampus [5,7–9], which is considered an essential region for the consolidation of short-term memories into stable long-term memories [10–12], and is mostly affected in DS.

Research on engram formation, maintenance and recall has uncovered several mechanisms and processes that are of paramount importance for memory function. Even so, only few studies have examined how engrams might be altered in cognitive disorders that are associated with memory deficits [9,13,14]. In fact, most of the critical mechanisms that contribute to adequate engram formation and recall are altered in mouse models of DS. For instance, plasticity deficits including impaired hippocampal LTP [15,16], increased synaptic depression [17] and deficits in structural plasticity [18,19] have been widely described in Ts65Dn, a trisomic mouse model of DS. Moreover, the excitatory/inhibitory balance, which is indispensable for engram formation, is altered in Ts65Dn mice and is thought to contribute to DS cognitive deficits [16,20,21]. Upregulation of CREB levels is one of the mechanisms that enhances neuronal excitability at the time of memory acquisition and its active form (p-CREB) is less abundant in the Ts65Dn hippocampus [22]. Similarly, *Kcnj6*, a gene that encodes for Kir3.2, an inwardly rectifying potassium channel, is triplicated in DS in Ts65Dn hippocampus. The upregulation of this gene has been associated with decreased excitability and is sufficient to impair both synaptic plasticity and memory [23]. Decreased neuronal activation was reported in the CA1 region of Ts65Dn during the exploration of novel environments [24]. Similarly, a reduction of Arc expression in the hippocampus after a novel object recognition test (NORT) was recently described in trisomic mice, compared to wild type (WT) littermates [25]. Altogether, this body of mounting evidence supports that engrams are probably defective in DS.

Here, we have explored whether engrams can form and reactivate in the Ts65Dn hippocampus. The hippocampus plays a critical role in the establishment of the contextual component of fear memories [26,27]. Specifically, the DG is involved in the discrimination between similar contexts [28]. In individuals with DS [29,30] and in mouse models of DS [31] the hippocampus is one of the most affected brain regions leading to hippocampal-dependent memories impairment [15,32]. We have tagged and manipulated engram cells and delineated the trisomic engram-specific alterations in the dorsal DG. Then, we have tested several strategies to revert engram-specific alterations to recover memory alterations in Ts65Dn mice.

## Results

### Increased sparsity of behaviorally induced neuronal activity upon learning in Ts65Dn in the Dentate Gyrus (DG)

Cognitive deficits have been extensively described in Ts65Dn mice, especially deficits in hippocampal-dependent tasks such as NORT [25,32] and CFC [15,33]. During the training session, Ts65Dn mice showed normal performance along the training session, with a significant increase of freezing during the training session to levels similar to the WT (ANOVA repeated measures; F(3,18) = 0.88; genotype effect N.S.; Figure 1BC). We next studied the pattern of neuronal activation during memory acquisition. We quantified the expression of c-Fos (Figure 1D) and Arc (Figure 1F) 90 min after learning. These immediate early genes (IEG) are transcriptionally induced by patterned synaptic activity and may reveal alterations in neuronal assembly recruitment upon learning. We focused on the DG since it is the input hippocampal gate and is important for pattern separation [34], crucial in contextual fear-conditioning paradigms [35]. In nontrained controls (home cage), the number of c-Fos and Arc positive cells was similar between genotypes (Post-hoc Tukey HSD; N.S., Figure 1E; N.S.; Figure 1G). However, upon learning, Ts65Dn mice displayed a significantly reduced number of c-Fos positive cells (Post-hoc Tukey HSD; p = 0.005; Figure 1E) and Arc positive cells (Post-hoc Tukey HSD; p = 0.007 Figure 1FG) in the DG compared to WT, indicating sparser neuronal activation upon learning.

**Figure 1.**
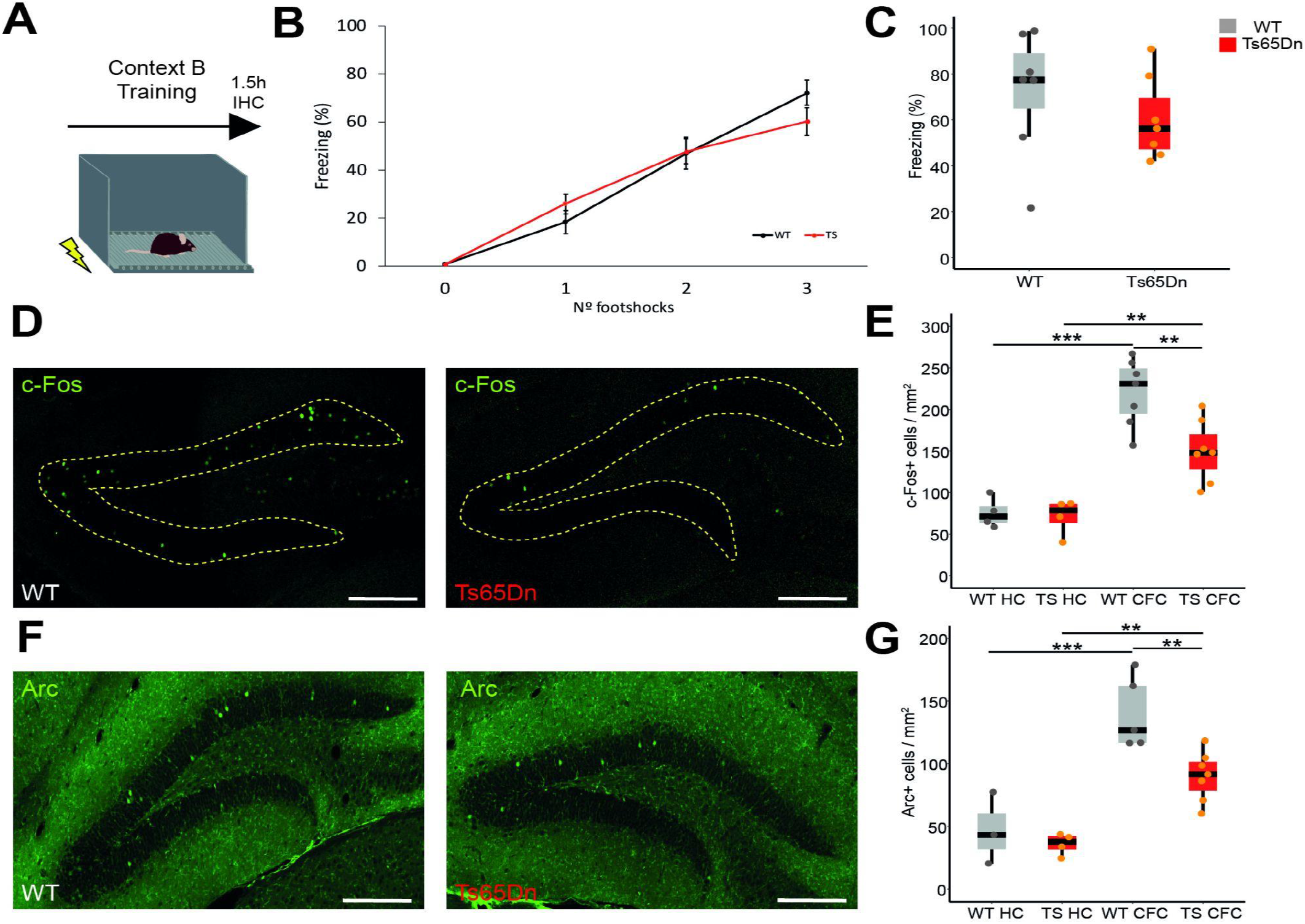
Sparse neuronal activation immediately after learning might contribute to defective engram allocation in Ts65Dn mice. **(A)** Behavioral schedule. Mice were trained in Context B. After 90 min, mice were sacrificed and the brain was extracted for immunohistochemistry. **(B)** Freezing behavior during the training session of contextual fear conditioning (CFC). Freezing was quantified before shock delivery (T 0), and after each shock (WT = 7 mice, Ts65Dn = 7 mice). **(C)** Freezing behavior after the last shock. **(D)** Representative images showing c-Fos expression (green) in the dorsal DG. Yellow dashed lines delineate the granule cell layer of the DG. Scale bars = 100 μm. **(E)** Quantification of c-Fos positive cells in the DG in non-trained (home cage; HC) controls (HC) and in mice submitted to CFC (WT HC = 4 mice, Ts65Dn HC = 4 mice, WT CFC = 7 mice, Ts65Dn CFC = 7 mice). **(F)** Representative images showing Arc expression (green) in the dorsal DG. Scale bars represent 100 μm. **(G)** Quantification of Arc positive in the DG in HC controls and in mice that underwent CFC. (WT HC = 3 mice, Ts65Dn HC = 4 mice, WT CFC = 5 mice, Ts65Dn CFC = 7 mice). Two-way ANOVA, TukeyHSD as post hoc. On the boxplots, the horizontal line indicates the median, the box indicates the first to third quartile of expression and whiskers indicate 1.5 × the interquartile range. \**P* < 0.05, ** *P* < 0.01, *** *P* < 0.001.

### Reduced engram size in Ts65Dn mice

The reduction in the number of active neurons upon CFC training in Ts65Dn mice suggested that engram size was reduced. However, the actual engram cells are those activated during learning and reactivated during recall, so, we used an activity-dependent engram tagging viral system (AAV_9_-cFos-tTA and AAV_9_-TRE_tight_-hM3Dq-mCherry) [36,37]. Temporal control over the activity-dependent expression of hM3Dq-mCherry was achieved by the removal of the doxycycline (DOX) diet (Figure 2A; see Methods). With this system, those cells activated during learning in the CFC training session were labeled with hM3Dq-mCherry, and memory was tested 24h after neuronal ensemble tagging (Figure 2B). Cells activated during learning were tagged with mCherry only in non-DOX conditions (Figure 2C and D). After the training session, constant DOX administration was reinitiated to prevent non-specific neuronal labeling. In the test session, Ts65Dn mice showed a significant reduction of freezing behavior compared to WT (Two-tailed T test; p < 0.001; Figure 2E) indicating a defective fear memory. To determine the actual engram size, brains were collected 90 min after memory recall and processed for IEG immunofluorescence to label memory recall activated cells (Figure 2F). In this experiment, the number of c-Fos+ cells (activated upon test) were significantly reduced in Ts65Dn mice (Two-tailed t test; p = 0.032; Figure 2G). However, as engram cells are those activated during learning and reactivated during memory recall, we quantified the number of c-Fos+ cells that co-expressed hM3Dq-mCherry. The number of engram cells in Ts65Dn were reduced compared to WT mice (Two-tailed T test; p = 0.017; Figure 2H), indicating reduced engram size in Ts65Dn mice. We also found a weak positive correlation between the freezing behavior and the number of engram cells, although it did not reach statistical significance (Spearman; Figure 2I).

**Figure 2.**
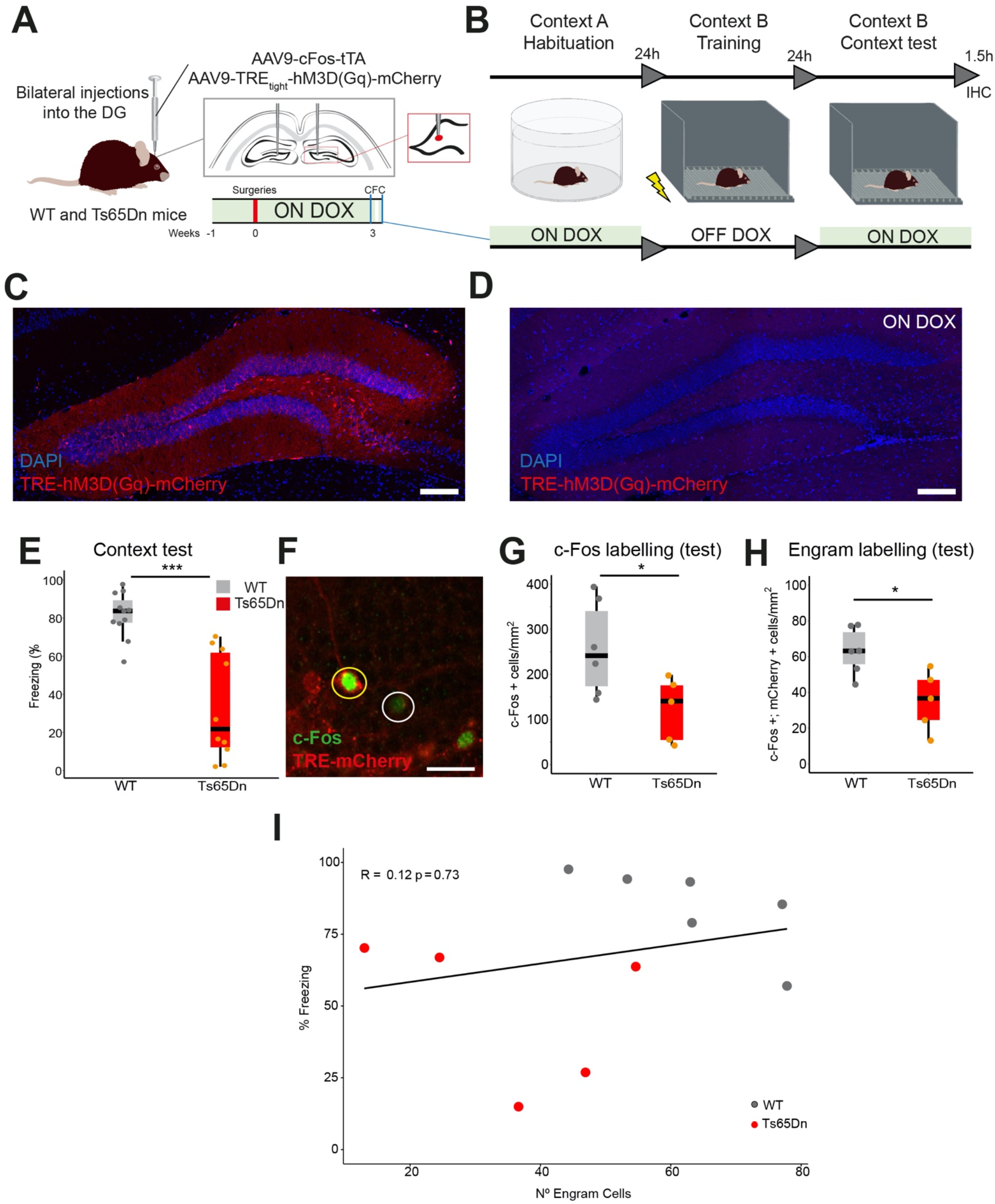
Reduced engram size and fear memory deficits in Ts65Dn mice. **(A)** AAV_9_-cFos-tTA and AAV_9_-TREtight-hM3D(Gq)-mCherry were injected into the dorsal DG of WT and Ts65Dn mice. **(B)** Mice were chronically ON DOX including the habituation to the neutral context (Context A). After habituation, DOX was removed and mice were trained in Context B 24h after. Then the DOX diet was resumed and mice were tested in Context B after 24h. Brains were extracted after 1.5h and processed for IEG quantification. **(C)** Representative image of mCherry+ cells in a DG section 24h after engram-tagging (OFF DOX). Scale bar = 100 μm. **(D)** Representative image of DG section keeping DOX diet (ON DOX) during the whole experiment, 24h after engram-tagging procedure. As expected, no mCherry+ cells are detected. Scale bar = 100 μm. **(E)** Memory recall in Context B. Ts65Dn mice froze significantly less than WT mice (WT saline = 11, Ts65Dn saline = 10). *P* < 0.001, Two-tailed T test. **(F)** c-Fos (green) and mCherry expressing cells 1.5h after memory recall test in the DG. **(G)** Density (cell/mm^2^) of c-Fos cells in the DG 1.5h after memory recall test (WT = 6, Ts65Dn = 5). *P* < 0.05, Two-tailed T test. **(H)** Density (cell/mm2) of double c-Fos+/mCherry+ cells in the DG 1.5h after memory recall test (WT = 6, Ts65Dn = 5). *P* < 0.05, Two-tailed T test. **(I)** Correlation between the number of engram (c-Fos+/mCherry+) cells and freezing behavior showing WT (black) and Ts65Dn (red). Spearman; R = 0,12, *p* = 0.73. On the boxplots, the horizontal line indicates the median, the box indicates the first to third quartile of expression and whiskers indicate 1.5 × the interquartile range. \**P* < 0.05, ** *P* < 0.01, *** *P* < 0.001

One important question when studying engram cells is their sufficiency and necessity to retrieve a specific memory [38]. To demonstrate necessity, in WT mice we utilized the engram tagging protocol with the inhibitory DREADD construct, AAV_9_-TREtight-hM4Di-mCherry (Supplementary Figure 1A). With this strategy, we were able to tag the learning-activated cells and impede their reactivation during memory recall using CNO. Engram cells in WT mice were tagged with hM4Di during the training session and 24 h after, CNO (3 mg/kg) was administered before re-exposure to the trained context (Supplementary Figure 1BC). We observed a significant reduction in freezing levels in WT animals treated with CNO compared to saline injected controls (Two-tailed T test; p = 0.0107; Supplementary Figure 1D), that would indicate that engram cells are necessary for proper memory retrieval.

To demonstrate sufficiency, we used the activity-dependent tagging protocol (AAV_9_-cFos-tTA and AAV_9_-TRE_tight_-hM3Dq-mCherry) to tag with mCherry and insert the excitatory DREADD (hM3Dq-mCherry) in those cells activated during acquisition (Supplementary Figure 1E). In this case, 24h after training, mice were administered (i.p.) with CNO (1 mg/kg) and 30 min later were exposed to the neutral Context A, in which mice did not receive the shocks (Supplementary Figure 1F). The expression of hM3Dq-mCherry was restricted to the DG (Supplementary Figure 1G).

In the CFC test, the neutral context (context A) did not produce freezing, as expected (Two-tailed T test; N.S.; Supplementary Figure 1H). By administering CNO, both WT and Ts65Dn mice showed freezing levels comparable to the ones observed in the conditioned context (B), indicating the successful activation of the fear memory engram. However, Ts65Dn mice froze significantly less than WT mice (Two-tailed T test; p = 0.04; Supplementary Figure 1I), indicating that, even in a neutral context, the activation of engram cells is sufficient to elicit memory recall in WT but, to a lesser extent in Ts65Dn, suggesting that the reduced engram size detected in trisomic mice might contribute to memory deficits.

Finally, since adequate engram reactivation is of paramount importance for memory recall, we tagged learning activated cells with hM3Dq-mCherry and 24 h later, we activated the engram cells with CNO (1 mg/kg) 30 min before re-exposing mice to the conditioned-context (Supplementary Figure 1J). Compared to the saline-injected group, the artificial activation of engram cells by CNO showed similar freezing values in WT and Ts65Dn mice (Pairwise Wilcox comparisons; p = N.S.; Supplementary Figure 1K), but artificial engram cell activation was not sufficient to overcome the memory deficits in trisomic mice (Pairwise Wilcox comparisons; p = 0.02; Supplementary Figure 1K).

### Trisomic engram cells do not display enhanced excitability after memory recall

Engram cells have been reported to undergo synaptic potentiation [7] after memory acquisition and their excitability is transiently enhanced after natural retrieval [8]. Because both synaptic potentiation [15,16] and neuronal excitability [39,40] are known to be impaired in Ts65Dn mice, we investigated whether trisomic engram cells are potentiated and increase their excitability to levels comparable to WT engram cells. Previous studies described a transitory increase in engram excitability 5 min after memory recall that remained until 3 h [8]. Since the neurons activated during the training session were tagged with hM3Dq-mCherry, we could identify those fluorescent cells in the DG. Mice were sacrificed mice 5 min after memory recall [8] to examine the electrophysiological profile of engram (hM3Dq-mCherry+) and non-engram (hM3Dq-mCherry-) cells by *ex vivo* whole-cell patch clamp recordings in the DG (Supplementary Figure 2AB). We found that action potential (AP) threshold was similar comparing hM3Dq-mCherry- and hM3Dq-mCherry+ both in WT and in Ts65Dn mice (Two-tailed t test; p = N.S.; Supplementary Figure 2A). Resting membrane potential was also similar between engram and non-engram cells in WT mice (Twotailed t test; p = N.S.; Supplementary Figure 2B), but it was higher in trisomic engram cells respect to non-engram cells (Two-tailed T test; p = 0.03; Supplementary Figure 2B). Membrane resistance showed a nonsignificant increase (Mann-Whitney; p = 0.079; Figure 3C, E) in WT engram cells compared to non-engram cells. We also found reduced rheobase in WT engram cells compared to non-engram cells (Mann-Whitney; p = 0.012; Figure 3C, F). However, in trisomic engram cells, we did not detect differences either in the membrane resistance or in the rheobase (Two-tailed t test; p = N.S.; Figure 3D, E-F). Interestingly, when plotting the rheobase against membrane resistance we could clearly see a higher mean difference between engram and non-engram cells in WT but not in Ts65Dn (Supplementary Figure 2C). These results indicate that WT engram cells enhance their excitability during memory recall. However, this was not the case of trisomic engram cells. We did not detect differences in the firing rate of both engram and non-engram cells either in WT (Supplementary Figure 3AB) or in Ts65Dn mice (ANOVA repeated measures = N.S.; Supplementary Figure 3CD).

**Figure 3.**
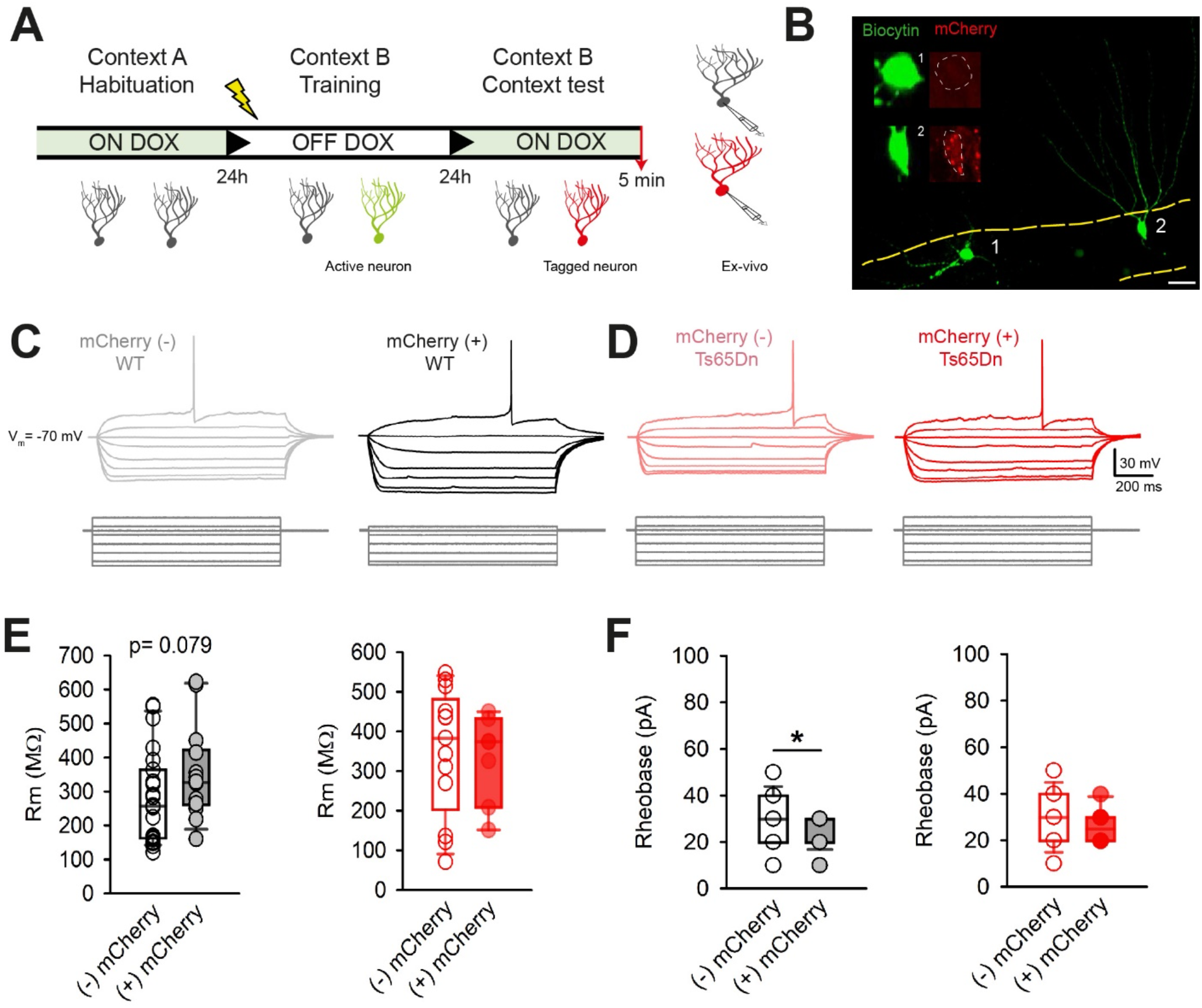
Electrophysiological profiling of DG engram cells. **(A)** Mice injected with the AAVs were habituated to the neutral Context A while ON DOX. After habituation, DOX was removed and mice were trained in Context B, 24 h after to allow tagging of active cells (in green). Then the DOX diet was resumed and mice were tested again in Context B after 24 h. Brains were extracted and processed for electrophysiological studies. **(B)** mCherry expressing cells (hM3Dq-mCherry+), were considered as engram cells while non-mCherry expressing cells (hM3Dq-mCherry-) as non-engram cells. Microphotograph showing non-engram (1) and engram (2) cells in the DG granule cell layer. Scale bar = 30 μm. **(C)** Representative traces of the first action potential elicited by increasing depolarizing pulses in non-engram (gray) and engram (black) in WT mice. **(D)** Representative traces of the first action potential elicited by increasing depolarizing pulses in non-engram (pink) and engram (red) in Ts65Dn mice. **(F)** Left: membrane resistance (Rm) of non-engram (black non-filled dots) and engram (gray filled dots) in WT mice. (WT non-engram = 23 cells from 8 mice, WT engram = 14 cells from 8 mice). Right: membrane resistance (Rm) of non-engram (red non-filled dots) and engram (red filled dots) in Ts65Dn mice. (Ts65Dn non-engram = 13 cells from 6 mice, Ts65Dn engram = 7 cells from 6 mice). **(H)** Left: rheobase of non-engram (black non-filled dots) and engram (gray filled dots) in WT mice. Right: rheobase of non-engram (red non-filled dots) and engram (red filled dots) in Ts65Dn mice. Mann Whitney. \**P* >0.05.

### Trisomic engram cells undergo structural plasticity

During memory consolidation the number of dendritic spines has been reported to increase in engram cells [7], and DS has been associated with abnormal dendritic spines [41]. Thus, we quantified the dendritic spines of engram and non-engram cells (hM3Dq-mCherry- and hM3Dq-mCherry+ neurons). Cells were filled with biocytin during electrophysiological recordings both in WT and in Ts65Dn mice (Figure 4AB). In accordance with previous reports, we found that spine density of hM3Dq-mCherry+ was significantly higher compared to hM3Dq-mCherry-cells (Post-hoc Tukey HSD; p < 0.001; Figure 4C) in WT mice. This increase was mainly accounted for by the mushroom and stubby spines (Post-hoc Pairwise Wilcoxon test; p < 0.001; Figure 4D). Trisomic hM3Dq-mCherry+ cells also exhibited an increased number of dendritic spines compared to hM3Dq-mCherry-neurons (Post-hoc Tukey HSD; p < 0.001; Figure 4D). There was a non-significant tendency to an increase in mushroom and stubby spines compared to trisomic hM3Dq-mCherry-cells (Post-hoc Pairwise Wilcoxon test; p = 0.056; Figure 4D), but the number of mature spines was significantly lower when compared to WT hM3Dq-mCherry+ neurons (Post-hoc Pairwise Wilcoxon test; p = 0.018; Figure 4D).

**Figure 4.**
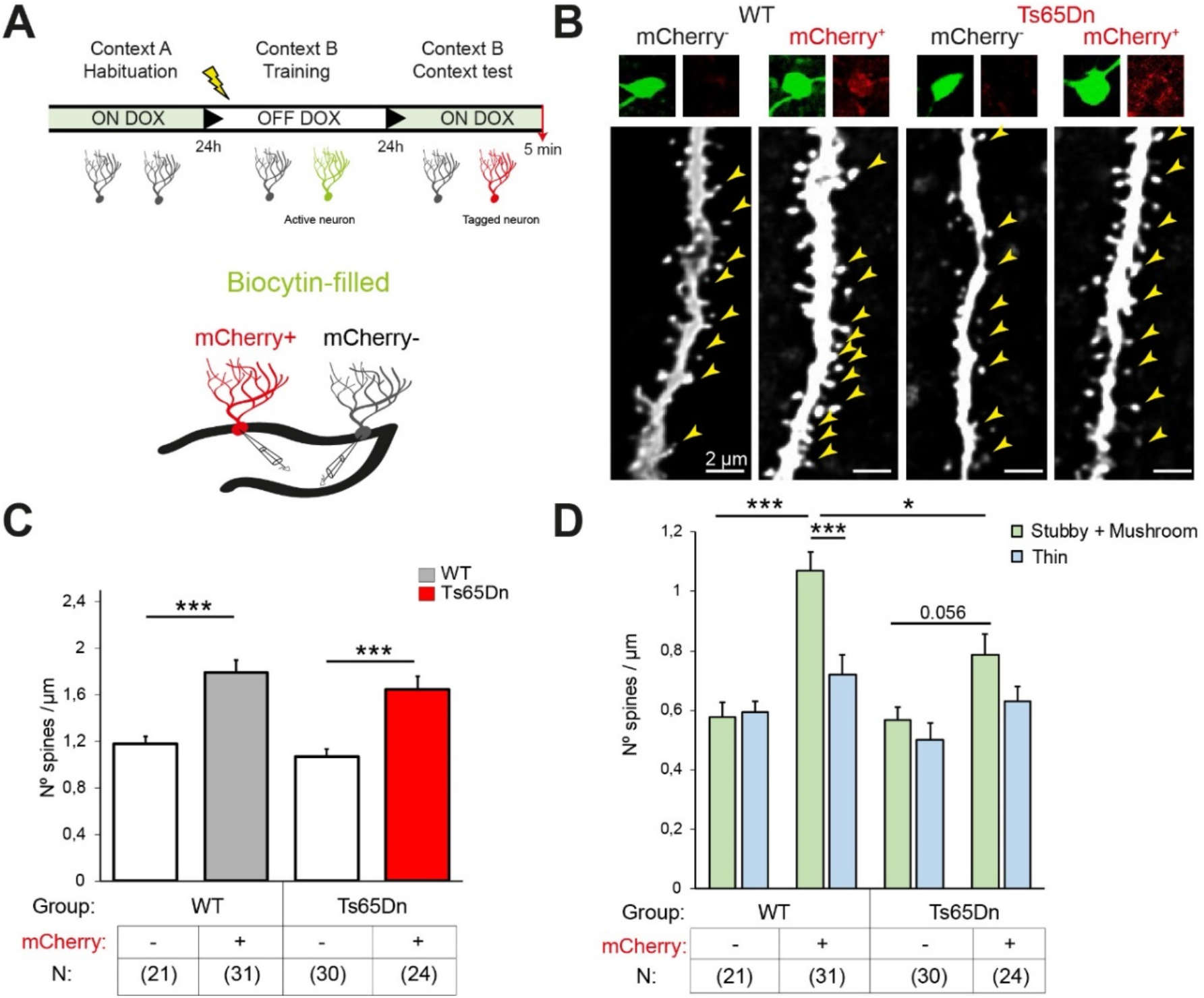
Trisomic engram cells undergo synaptic potentiation but present a fewer number of mature spines. **(A)** Mice injected with the AAVs were habituated in the neutral context (Context) while ON DOX. After habituation, DOX was removed and mice were trained in Context B 24 h after to allow tagging of active cells (in green). Then the DOX diet was resumed and mice were tested again in Context B after 24 h. Brains were extracted and processed for electrophysiological studies. During electrophysiological recording, granule cells were filled with biocytin for posterior reconstruction. **(B)** Representative images of mCherry- and mCherry+ cells both in WT and in Ts65Dn mice (arrow heads: dendritic spines). Scale bar = 2 μm. **(C)** Average dendritic spine density showing an increase in mCherry+ cells (WT non-engram = 21 dendritic fragments from 6 cells, WT engram = 31 dendritic fragments from 6 cells, Ts65Dn non-engram = 30 dendritic fragments from 7 cells, Ts65Dn engram = 24 dendritic fragments from 5 cells). Two-way ANOVA, Tukey HSD as post hoc. **(D)** Average dendritic spine density by spine type. (WT non-engram = 21 dendritic fragments from 6 cells, WT engram = 31 dendritic fragments from 6 cells, Ts65Dn non-engram = 30 dendritic fragments from 7 cells, Ts65Dn engram = 24 dendritic fragments from 5 cells). Two-way ANOVA, Tukey HSD as post hoc. \**P* < 0.05, ** *P* < 0.01. Data are expressed as mean ± SEM.

### Overexpression of CREB in the DG does not rescue engram size and memory deficits in Ts65Dn mice

Several lines of evidence suggested that CREB plays a key role in neuronal ensemble allocation to an engram [6], and we previously showed that the levels of phosphorylated CREB was reduced in Ts65Dn mice [22]. In the present experiments, we confirmed that the levels of CREB in granule cells were reduced in Ts65Dn after the CFC training session compared to WT mice (Twotailed T test; p < 0.001; Figure 5AB). Thus, we decided to upregulate CREB expression in the DG prior to the acquisition of contextual fear conditioning memory to determine whether we could rescue engram size and memory in Ts65Dn. We injected AAV_2_-hSyn-CREB-EGFP into the dorsal DG 3 weeks before CFC training (Figure 5C). CREB-EGFP expression was restricted to the granule cell layer of the dorsal hippocampus (Figure 5D). However, we could not confirm previous observations [42], as in our experiments despite the upregulation of CREB expression WT mice did not increase freezing levels, indicating no effect on memory. Similarly, overexpression of CREB did not enhance memory in Ts65Dn mice (Two-tailed T test; p = 0.0273; Figure 5EF). Indeed, the number of c-Fos+ neurons during the context test was reduced in Ts65Dn compared to WT mice (Two-tailed T test; p = 0.0277; Figure 5G) even though AAV-CREB-EGFP was successful in elevating CREB expression levels in neurons expressing CREB-EGFP both in WT and in Ts65Dn mice (Post-hoc Tukey HSD; p < 0.001; Figure 5HI).

**Figure 5.**
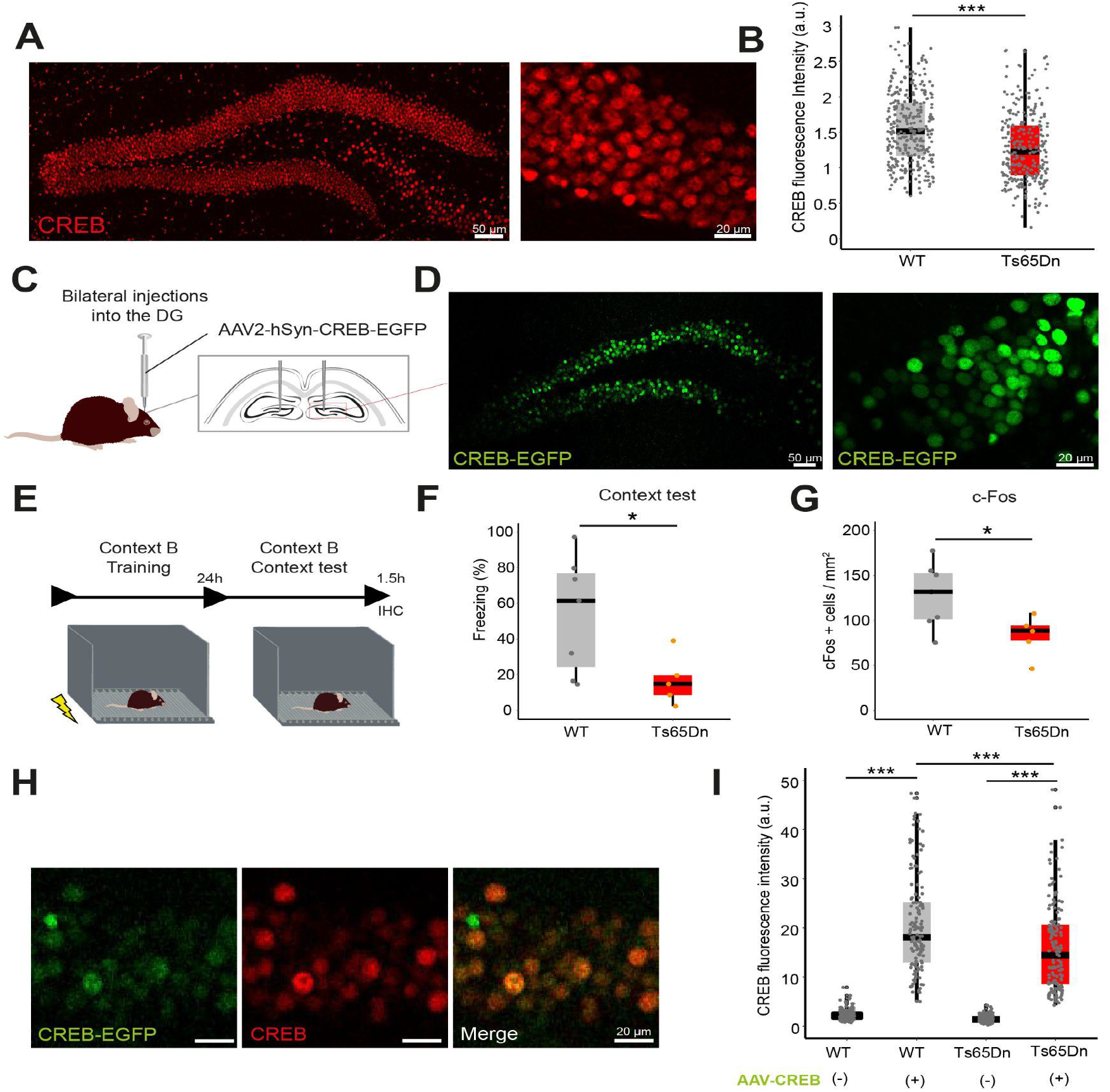
CREB overexpression does not rescue memory deficits in Ts65Dn mice. **(A)** Left: representative images showing CREB expression in the dorsal DG. Scale bar = 50 μm. Right: Image magnification showing CREB expression. Scale bar = 20 μm. **(B)** Fluorescence intensity granule cells in the DG in arbitrary units (WT = 328 granule cells from 6 mice, Ts65Dn = 280 granule cells from 7 mice). Two-tailed T test. **(C)** AAV_2_-CREB-EGFP was injected into the DG of WT and Ts65Dn mice. **(D)** Left: Representative image showing CREB-EGFP expression in the dorsal DG. Scale bar = 50 μm. Right: Magnification showing the pattern of CREB-EGFP expression. Scale bar = 20 μm. **(E)** Experimental schedule of CFC. **(F)** Percentage of freezing of WT (gray) and Ts65Dn (red) in the context B during the test session (WT = 7, Ts65Dn = 5). **(G)** c-Fos density in the DG 1.5h after memory recall test (WT = 7, Ts65Dn = 5). Two-tailed T test. **(H)** Representative image showing expression of AAV-CREB-EGFP (green), CREB (red) and the merged image in the granule cell layer of the hippocampus. Scale bar = 20 μm. **(I)** CREB fluorescence intensity on CREB-EGFP negative and positive cells both in WT and Ts65Dn mice (WT AAV-CREB negative = 158 cells from 3 mice, WT AAV-CREB positive = 149 cells from 3 mice, Ts65Dn AAV-CREB negative = 143 cells from 3 mice, Ts65Dn AAV-CREB positive = 149 cells from 3 mice). Two-way ANOVA, Tukey HSD as Post-hoc. On the boxplots, the horizontal line indicates the median, the box indicates the first to third quartile of expression and whiskers indicate 1.5 × the interquartile range. *** *P* < 0.001, * *p* < 0.05.

## Discussion

We here evaluated engram formation and reactivation in the context of DS. Our hypothesis was that DS is a model of inherently defective engram formation and reactivation. Previous work in DS mouse models have showed that memory deficits may arise from a failure to convert transitory learning-induced neuronal activity into longterm memory traces [41]. Thus, we first evaluated whether the pattern of neuronal activation immediately after learning was impaired in Ts65Dn using a CFC paradigm [15]. Ts65Dn mice showed training-induced freezing levels similar to WT mice, indicating no impairment in CFC learning. In agreement with our findings, other groups also reported no differences in the freezing behavior during the training session in Ts65Dn mice [43,44]. However, even though Ts65Dn mice were able to learn, the number of learning-activated neurons (c-Fos+ and Arc+ cells) were reduced in Ts65Dn DG, indicating that sparser learning-induced neuronal ensemble activity is not accompanied by learning deficits in Ts65Dn. Similar results were obtained in a mouse model of AD, in which, even though training-induced freezing levels were similar to WT mice, the number of activated neurons were reduced during training [9]. Our findings suggest that training-induced freezing does not predict the number of active neurons during memory acquisition and that alternative mechanisms might explain the reduced number of active neurons in Ts65Dn mice. A sparser neuronal ensemble activation was also detected in a previous study in the CA1 region of Ts65Dn, upon exposure to novel environments [24]. In addition, a recent study from our laboratory found a lack of upregulation of Arc levels immediately after learning in Ts65Dn mice while Arc levels did not differ in basal conditions [25]. It might be possible that cell-autonomous mechanisms, such as impaired intrinsic excitability [39,40,45] or non-autonomous factors might lower the number of active neurons in response to learning cues. Although we did not directly evaluate neuronal activity after learning, the reduced number of granule cells expressing c-Fos and Arc in Ts65Dn mice would be in line with a reduced neuronal activity. In fact, studies on animal models of AD also observed sparser neuronal activation during learning [9] along with plasticity defects [9,14] and engram size reduction [9,13].

Thus, and even though c-Fos is not a direct marker of the engram, our findings could be interpreted as a deficit in the allocation of stable and long-lasting neuronal ensembles during learning, as this depends on IEGs production [46,47]. To proof this assumption, we next used activity-dependent tagging with the fluorescent reporter mCherry to identify the neurons that were activated both during the acquisition of a contextual fear memory and reactivated during memory recall (c-Fos and mCherry expressing neurons), e.g. engram cells. Remarkably, the number of engram cells was significantly reduced in the Ts65Dn granule cell layer, and correlated with memory performance both in WT and in Ts65Dn mice.

Although neuronal ensemble activation during learning is assumed to be relevant for the correct representation of memories in the brain [46,48,49], it has been suggested that engram size is quite stereotyped and that deviations from the optimal size might prevent engram networks to reactivate memory. For instance, larger engram size could lead to better discrimination due to a greater cellular reactivation in the conditioned context [48]. Considering the DG has a role in memory resolution [50,51], the greater the neuronal ensembles recruited during learning would promote an improved memory specificity. As such, it could be assumed that the degree of learning and memory specificity corresponds with neuronal activity in a subset of DG cells that were active during both learning and recall [48].

Reduced engram size, on the other hand, would make it more difficult to distinguish between comparable contexts since less contextual information would be encoded into the same the memory trace, resulting in memory instability and poorer discrimination [48]. In fact, Ts65Dn are impaired in a pattern separation task [52]. Although in our experiments engram size was not directly related to learning capability, it is commonly accepted that accessing the memory engrams is necessary for memory recall [53], and thus, engram size in the trisomic mice would affect recall. Indeed, engram studies on AD and in FXS also observed reduced engram size upon memory retrieval [9,13,14]. However, while in AD mouse models, engram size was restored and memory could be recalled by artificial reactivation of the engram using optogenetics, in the FXS model, memory deficits and impaired engram reactivation were rescued by promoting neuronal plasticity before learning by submitting mice to EE [13]. Based on these studies, we could speculate that impaired trisomic memory engram reactivation may be due to i) defective engram allocation, as suggested by the reduced c-Fos expression, or ii) impaired engram reactivation, as indicated by the reduced proportion of mCherry/c-Fos+ cells. In order to decipher this question, we decided to activate the engram neurons that were tagged with mCherry and expressed an excitatory DREADD (hM3Dq-mCherry) in the trained context using CNO (1 mg/kg). However, and contrary to what was observed in the AD scenario, memory was not rescued in Ts65Dn. This result would suggest that memory deficits in Ts65Dn are not contributed by impaired engram reactivation since artificial engram activation by CNO does not rescue the reduced memory during natural recall in trisomic mice. Taken together, these results would indicate that reduced engram size underlie impaired contextual fear memory recall in Ts65Dn mice.

Besides the number of engram cells, other critical factors such as neuronal excitability and connectivity have been found to be important for memory retrieval. Alterations in the capability of engram cells to form a stable engram network and/or to properly reactivate under specific recall cues can hinder memories to be retrieved. For instance, a recent study found that engram cell excitability influences the efficacy of memory retrieval [8]. The authors found that recall cues induce a transient enhancement of engram cell excitability that leads to improved context recognition.

Thus, we next studied the intrinsic properties of engram and non-engram cells to detect possible alterations in neuronal excitability and neuronal plasticity [5,7,9]. By means of whole-cell patch clamp recordings, we were able to determine the passive and active electrical properties of both WT and trisomic mCherry- and mCherry+ cells, as a proxy of non-engram and engram cells. In WT mice, we confirmed the reduced rheobase in mCherry+ compared to mCherry-cells that Pignatelli et al. found in engram cells immediately after memory retrieval [8]. This enhancement of neuronal excitability of mCherry+ cells was not observed in Ts65Dn reinforcing the concept that reduced excitability might prevent memory to be entirely recalled.

It is well known that CREB-mediated mechanisms increase excitability by reducing voltage-gated K^+^ currents [54,55]. In fact, CREB upregulation enhances cell excitability [56], which is crucial in the engram allocation process. Previously in the lab, we showed altered levels of p-CREB in the Ts65Dn hippocampus [22]. Thus, we explored whether alterations in CREB signaling could account for the smaller engram size in Ts65Dn mice. We found that in Ts65Dn, CREB levels were reduced after memory acquisition in the DG compared to WT littermates. In light of the key role of CREB on the engram allocation process, and given its reduced levels in trisomic mice, we decided to upregulate its expression in the DG using a custom-made AAV_2_-hSyn-CREB-EGFP. CREB-EGFP was expressed in granule cells and was effective to upregulate CREB expression both in WT and Ts65Dn mice. However, neither in WT mice nor in Ts65Dn, CREB upregulation was able to improve memory performance or neuronal activation after memory recall. Our results may seem contradictory to previous work showing that increasing the function of CREB in principal lateral amygdala neurons increased the likelihood to become part of a fear memory engram [6]. It could be argued that WT mice already perform at very high levels, thus possibly indicating a ceiling effect. However, a result similar to our finding was observed in a previous study, in which CREB upregulation did not enhance neuronal activation nor memory performance in WT mice [14]. Importantly, in that work it increased the number of active neurons and rescued memory deficits in a mouse model of AD [14]. Instead, we show that this is not the case in Ts65Dn mice, in which the upregulation of CREB alone might not be sufficient to increase the number of active neurons suitable for being allocated to an engram at the time of learning. As our construct CREB-EGFP was under the hSyn promoter, CREB was upregulated in interneurons, so it is also conceivable that elevated interneuron excitability could lead to a greater inhibition of excitatory trisomic granule cells. In order to exclude this possibility, it would be interesting to investigate whether enhancing CREB expression exclusively in excitatory granule cells might be effective to overcome sparser neuronal activation in Ts65Dn mice. It is important to bear in mind that the connectivity of neural subnetworks activated by learning depends on the strength and numbers of synapses between neurons, which influences the neuronal activation and, thereby, determines the way engrams stabilize and reactivate. Engram cells undergo structural and functional plasticity [7–9]. Previous studies on retrograde amnesia [7] and AD [9] have described that the lack of memory retrieval originates from a disruption of the consolidation phase that occurs during the transition from a fragile and recently encoded memory engram to a stable and mature memory engram. During this consolidation process, engram cells persistently increase their synaptic strength and spine density. It was thus important given the altered structural plasticity detected in trisomic mice [41] to study dendritic spine density of engram and non-engram cells in WT and in Ts65Dn mice. In accordance with previous results, WT spine density was increased in engram compared to non-engram cells [7,9]. However, spine density was also slightly, although not significantly enhanced in trisomic engram cells. Interestingly, in WT engram, cells the increased spine number was contributed by mature spines, in both genotypes. Even so, the number of mature spines was higher in WT engram cells compared to Ts65Dn engram cells. Our results do not confirm previous findings of reduced number of spines in the trisomic DG [18,19,57,58], as we did not detect differences in spine density between WT and Ts65Dn granule cells. It is possible that differences in the quantification methods and the age of the animals might account for these variations. For instance, some studies were performed at P15 [18,19] while others at 6 months-old Ts65Dn mice [57]. Moreover, most of these studies used Golgi staining [18,19,58] or electron microscopy [57] that might allow a higher resolution of spines and, therefore, to detect smaller differences in spine number. In our case, the method used to identify engram (mCherry+) cells using stereotaxic injection of the activity-dependent tagging system along with the whole-cell patch clamp recordings, might add some degree of variability that might prevent us from detecting differences in spine number in granule cells. Even so, we were able to reproduce previous results of increased spinogenesis in WT engram cells [7].

Again, the trisomic case is different from both anterograde amnesia and AD. Whilst in anterograde amnesia and AD, structural plasticity is not detected in engram cells [7,9], Ts65Dn engram cells showed a slight increase in dendritic spine density. These results suggest that, as shown in Ts65Dn mice, the reduced structural plasticity is not sufficient to support engram stabilization. This finding may relate to the reduced neuronal ensemble activation during memory acquisition, as one would expect reduced activity to impair structural plasticity.

In conclusion, to the best of our knowledge, herein, we have performed the first characterization of the trisomic engram cells in the hippocampus, thereby identifying key cellular and molecular changes that might lead to defective engrams in DS. The identification of engram dysfunction in DS along with several candidate mechanisms will open new research avenues to explore DS from a completely different perspective.

## Author contributions

MD and AFB conceived the study, designed and coordinated the study and wrote the manuscript. AFB and MS conducted *in vivo* animal experiments. AFB analyzed histological preparations. AZM designed and conducted the electrophysiology experiments. All authors revised and corrected the final version of the manuscript.

## Materials and Methods

### Animals

Ts(17^16^)65Dn (Ts65Dn) mice were obtained through crossings of a B6EiC3Sn a/A-Ts (17^16^)65Dn (Ts65Dn) female to B6C3F1/J males purchased from The Jackson Laboratory (Bar Harbor, USA). Genotyping was performed by amplifying genomic DNA obtained from the mice tail as described in (Liu et al., 2003). Mice had access to food and water *ad libitum* in controlled laboratory conditions with temperature maintained at 22 ± 1°C and humidity at 55 ± 10% on a 12h light/dark cycle (lights off 20:00 h). Mice were socially housed in numbers of two to four littermates. The colony of Ts65Dn mice was maintained in the Animal Facilities in the Barcelona Biomedical Research Park (PRBB, Barcelona, Spain).

According to Directive 63/2010 and Member States’ implementation of it, all trials followed the “Three Rs” principle of replacement, reduction, and refinement. The investigation was conducted in accordance with the Standards for Use of Laboratory Animals No. A5388-01 (NIH) and local (Law 32/2007) and European regulations as well as MDS 0040P2 and the Ethics Committee of Parc de Recerca Biomèdica (Comité Ético de Experimentación Animal del PRBB (CEEA-PRBB)). A/ES/05/I-13 and A/ES/05/14 grant the CRG permission to work with genetically modified organisms. See the Ethics section for further information.

### Experimental strategy

We have investigated engram-specific alterations using engram tagging techniques and chemogenetic intervention in Ts65Dn. In the first experiments using immunostaining and recombinant AAV_9_-TRE-tight-hM3Dq-mCherry, we characterized neuronal activation pattern in Ts65Dn in 3-month-old male mice both after learning and after recall. We quantified the number of cells expressing c-Fos and Arc IEGs in the dentate gyrus. For the behavioral experiments, all mice, unless otherwise specified, were males aged 10-12 weeks at the time of learning. For engram tagging and manipulation and CREB overexpression experiments, males were aged 8-10 weeks at the time of virus injection. For engram tagging experiments, mice treated with food containing 40 mg/kg doxycycline (DOX) at least 7 days before the surgery. After surgery, mice remained on DOX except for the engram tagging time window. For *ex vivo* whole-cell patch clamp electrophysiology experiments mice were 11-13 weeks old at the time of the experiment. Engram tagging and manipulation was performed using chemogenetic intervention with clozapine N-oxide (CNO) to determine whether the trisomic engram was not properly formed, or not properly reactivated.

### Engram tagging and manipulation techniques

#### Viral constructs

In order to detect and to manipulate memory engram cells, we used a construct in which the excitatory DREADD hM3D(Gq) or the inhibitory DREADD hM4D(Gi) were fused to mCherry under the control of the tetracycline-responsive element (AAV_9_-TRE-tight-hM3D(Gq)-mCherry). To allow the combination with AAV_9_-cFos-tTA into the targeted zone, the dorsal DG, we used a double viral system. For engram activation we combined (1:2) AAV_9_-TRE-tight-hM3D(Gq)-mCherry and AAV_9_-cFos-tTA. For engram inactivation we combined (1:2) AAV_9_-TRE-tight-hM4D(Gi)-mCherry (diluted 1:5 in PBS) and AAV_9_-cFos-tTA. pAAV-cFos-tTA-pA and pAAV-PTRE-tight-hM3Dq-mCherry were a gift from William Wisden (Addgene plasmid #66794 and #66795). AAV vectors were serotyped with AAV_9_ coat proteins and packaged at the Viral Production Unit (UPV) at the Universitat Autònoma de Barcelona. Viral titers were 1.6 × 10^13^ genome copy (GC)/mL for AAV_9_-cFos-tTA and AAV_9_-TREtight-hM3Dq-mCherry. AAV_9_-TRE_tight_-hM4Di-mCherry was generated by replacing the hM3Dq-mCherry of #66795 for hM4Di-mCherry from #50479 (Addgene plasmid, gift from Bryan Roth). Viral titer was 1 × 10^13^ genome copy (GC)/ mL. AAV_2_-hSyn-CREB-EGFP was constructed by substituting the CMV promoter for the hSyn from the AAV-CREB (Addgene plasmid #68550; gift from Eric Nestler), serotyped with AAV_2_ coat proteins and packaged at Viral Production Unit at the UPV. Viral titer was 8 × 10^12^ genome copy (GC)/ mL for AAV_2_-hSyn-CREB-EGFP.

### Pharmacogenetics (DREADD)

For the inhibition or reactivation of the DREADDS we used CNO, an inactive form of clozapine drug. The synthetic ligand CNO binds to the modified human muscarinic receptor hM3D(Gq) or hM4D(Gi). CNO was dissolved in DMSO and diluted in 0.9% saline to yield a final DMSO concentration of 0.5%. Control animals received 0.5% DMSO saline solutions. For engram activation studies, 1 mg/kg CNO was intraperitoneally injected 30 min before the behavioral assays. For engram inactivation experiment, 3 mg/kg CNO dose was used. None of the doses of CNO induced any behavioral alterations or signs of seizure activity.

### Stereotaxic surgery injection

Intracerebral injections in DG were performed bilaterally at Bregma −2.2 mm AP, ± 1.3 mm ML, −1.9 mm DV with the aid of a stereotaxic apparatus (Stoelting 51730). 8-10 weeks old male mice were anesthetized using ketamidol (7.5 mg/kg) and medetomidin (0.2 mg/kg). Fur was shaved from the incision site. Skin was wiped with ethanol 70% and small incisions were made along the midline to expose bregma and injection sites. Craniotomies were performed using a 0.45 mm diameter stereotaxic microdrill (RWD Life Science, model 78001). 150 nl of each virus (1:2) were injected per hemisphere at a rate of injection of 50 nl/min during 6 min through a 33-gauge cannula (Plastics One, C235I/Spc) attached to a Hamilton microsyringe (1701N; Hamilton) connected to a syringe pump (PHD 2000, Harvard Apparatus). Cannulas remained 10 min after injection to allow virus diffusion and were slowly withdrawn during 5 min. Skin was sutured and mice were treated with 0.03 mg/kg buprenorphine as analgesic. Mice were recovered from anesthesia by atipemazol (1 mg/kg) and maintained on a heating pad until fully recovered. Mice were allowed to recover for 3 weeks before experimentation to allow construct expression. After sacrifice, all injection sites were verified histologically. Only those mice in which virus expression was restricted to the dorsal DG were included in the analyses.

### Behavioral assays

#### Contextual fear-conditioning (CFC) paradigm

CFC is a hippocampal-dependent test used to interrogate associative learning and to study engram formation and reactivation. Importantly fear memory is altered in trisomic mice. We adapted a training paradigm for contextual fear-conditioning (3 shocks; 0.6 mA, separated by 60 s) used in previous activity-dependent tagging studies (Figure 1A) [5,7]. Briefly, mice are introduced into a new environment and an aversive stimulus is delivered at different time points. The next day mice are reexposed to the same context without any shocks delivered. If mice recall and associate the context to the aversive stimuli, they will typically exhibit a freezing reaction when placed back in that setting. As a reaction to fear, freezing is described as “lack of movement other than breathing.”

Two different contexts were employed in two different cages, one neutral not associated with fear-related learning (Context A) and a second one (Context B) that was paired to the unconditioned (shock) stimulus. Context A (neutral) consisted in a chamber (29 × 25 × 22 cm) with Perspex floors and transparent circular ceilings and Context B (30 × 25 × 33 cm) had grid floors, and opaque square ceilings. After each session, the apparatus was cleaned with 70% ethanol.

All mice were individually handled and habituated to the investigator during three days before the experiment. One day before conditioning (training) mice were habituated in Context A for 3 min. No shocks were delivered in this session. Immediately after the habituation, DOX diet was removed and mice switched to a standard diet for a period of 24h, to allow expression of the doxycycline sensitive tetracycline transactivator (tTA) under the control of the c-Fos promoter. After habituation mice were placed back in their home cages in the holding room.

After 24h, mice were trained in Context B during 300 s, with three 0.6 mA shocks of 2s duration delivered at 120 s, 180 s and 240 s, respectively [7]. After training, DOX diet administration was reestablished to restrict the expression of the construct to the learning period. After training mice were placed back to their home cages.

All testing sessions in Context B were 180 s in duration. Testing conditions were identical to training conditioning, except that no shocks were delivered. At the end of each session mice were placed in their home cages. Freezing behavior (>800 ms immobility) was automatically detected by Packwin 2.0 software (Panlab, Harvard Apparatus). Cages were calibrated according to manufacturer instructions each

### Histology

#### Immunohistochemistry

In order to quantify IEGs expression either after memory acquisition or memory recall, mice were transcardially perfused with ice-cold PBS followed by 4% paraformaldehyde (PFA) in PBS (pH 7.4). Brains were extracted and post-fixed in 4% PFA at 4°C overnight. Brains were then transferred to PBS and 40 μm coronal consecutive brain sections were obtained employing a vibratome (Leica VT1200S, Leica Microsystems), collected in PBS and stored in cryoprotective solution (40% PBS, 30%, glycerol and 30% polyethylene glycol) for long-term storage. For immunofluorescence studies, 4-6 sections per mice were selected centered on the injection sites and according to stereotaxic coordinates Bregma, −1.54 to −2.54 mm, (mouse brain atlas; Franklin & Paxinos, 2012) with the aid of a bright-field microscope (Zeiss Cell Observer HS; Zeiss). Brain sections were washed with PBS (3 × 10 min). Then, sections were permeabilized with 0.5 % Triton X-100 in PBS (PBS-T 0.5 %) (3 × 15 min) and blocked with 10% of Normal Goat Serum (NGS) for two h at room temperature (RT). Sections incubated in PBS-T 0.5% and NGS 5 % with the primary antibodies overnight at 4°C washed again (PBS-T 0.5 % 3×15 min) and incubated with the secondary antibodies (PBS-T 0.5 % + NGS 5 %) for two h at room temperature protected from light. Finally, samples were washed with PBS-T 0.5 % (3×15 min) followed by PBS washing (3×10 min) to remove the detergent and sections were mounted and coverslipped into a pre-cleaned glass slide with Fluoromount-G medium with DAPI (Thermo Fisher Scientific #00-4959-52). c-Fos was stained with rabbit anti-c-Fos (1:1000, Santacruz, #Sc-7202) and visualized with anti-rabbit Alexa-647 (1:500; Thermo Fisher Scientific, #A-21443). Arc was stained with mouse anti-Arc(C-7) (1:400; Santa Cruz, #sc-17839) and visualized with anti-mouse Alexa-488 (1:500; Thermo Fisher Scientific, #A-11001). CREB was stained with rabbit anti-CREB (1:1000; Millipore, #06863) and visualized with anti-mouse Alexa-555 (1:500; Thermo Fisher Scientific, #A-32732). Prior to immunostaining, an optimization of the primary antibodies and PBS-T conditions was performed. Serial dilutions of primary antibodies ranging from 1:100 to 1:1000 were prepared while maintaining the secondary antibody concentration constant (1:500). By confocal microscopy, the best primary antibody concentration was selected taking into account the achievement of low background noise and the signal level obtained with the same laser configuration.

#### Biocytin immunohistochemistry protocol after whole-cell patch clamp recording

To visualize and quantify the dendritic spines of engram and non-engram cells that were subjected to electrophysiological recordings, 300 μm slices were first fixed in 4% PFA for 24h. Brain slices were washed with Tris-Buffered Saline (TBS) (3 × 10 min). Then, sections were permeabilized with 0.3% Triton X100 in TBS (TBS-T) and blocked with 20% NGS for 1 h at RT. Then slices were incubated with Streptavidin, Alexa Fluor 488 conjugate (1:1000, Thermo Fisher; #S32357) in 0.3% TBS-T with 1% NGS overnight at 4°C. Samples were washed with TBS (3 × 10 min) and sections were mounted and coverslipped intro pre-cleaned glass slice with Fluoromount-G medium with DAPI (Thermo Fisher Scientific #00-4959-52).

#### Cell counting

In order to quantify the number of engram cells in the DG (c-Fos and hM3D(Gq)-mCherry-expressing cells), 40 μm coronal sections were taken from the dorsal hippocampus centered in the coordinates where the viruses were injected (−1,54 to −2,54 mm AP; relative to bregma). Cell densities are expressed as cells/mm^2^. Confocal fluorescence images were acquired on a Leica TCS SP5 inverted scanning laser microscope using a 20x/0.70 NA objective. Cell counting was performed using the Cell Counter plugin on ImageJ software (NIH, Bethesda) in a z-stack (3 μm step size). The somatic layer of the granule cell layer was selected as region of interest (ROI) and was manually delineated according to the DAPI signal in every section.

Alexa 488 and Alexa 568 channels were filtered and combined to produce composite images. Equal cutoff thresholds were applied to remove signal background from images. The number of double positive (hM3Dq-mCherry and c-Fos) and single positive (c-Fos) cells were counted in the DG in 3-6 consecutive coronal sections (spaced 200 μm between them) per mouse. The same procedure was used to quantify c-Fos, Arc, the only difference being the channels used to create composite images. Data was analyzed using R studio. Imaging and quantifications were performed blind to experimental conditions

#### Spine density analysis

Engram cells were labeled by the c-Fos-tTA-driven induction of hM3Dq-mCherry (see Viral Construct section). mCherry-expressing (engram) and mCherry non-expressing (non-engram) cells both in WT and in Ts65Dn DG cells were labeled with biocytin during whole-cell patch clamp recordings. Slices were fixed in 4% PFA for 24 h and immunofluorescence protocol against biocytin was performed. mCherry signal was also amplified using immunohistochemistry procedures. Fluorescence Z-stacks of dendritic spines were taken by confocal microscopy (Leica SP5 inverted, Leica Microsystems), using 63x glycerol immersion objective. Images were deconvoluted using Huygens essential software. To minimize quenching of fluorescence, z-stacks were rapidly scanned at 0.2 μm increments. We only included dendrites for analysis if the labeling was bright and continuous throughout its course. Primary dendrites were not included in the analysis. All images were processed in batch using the same template (available upon request). Deconvoluted images were imported into NeuronStudio and were semi-automatically analyzed blind to experimental conditions. 5-7 granule cells were analyzed for dendritic spine quantification (n = 5-7 cells per group; n = 4 groups). For spine density, we analyzed different fragments of each cell quantifying at least 100 μm of secondary dendritic spines. All dendritic spines were at least 50 μm apart from the neuronal soma. Primary dendrites were not counted. The density of spines was calculated as the number of spines in every dendritic fragment’s length.

#### CREB fluorescence intensity measurements

In order to quantify the fluorescence intensity of CREB in euploid and trisomic granule neurons, confocal fluorescence images were acquired at 20x magnification on a Leica TCS SP5 inverted scanning laser microscope (Leica Microsystems) creating a composite image of the entire dorsal hippocampus at 16 bits. Confocal acquisition settings were maintained constant for all the samples and all images were taken the same day. Tissue was only exposed to the lasers during the moment of image acquisition to prevent photobleaching. The mean intensity of granule cell somas was performed by manually delineating the somas in accordance with the CREB signal using ImageJ software (NIH, Bethesda). Signal background was subtracted for every region and image.

### Electrophysiology

#### Ex vivo whole-cell patch clamp recordings

Engram was tagged as previously mentioned and mice were decapitated 5 min after natural recall in Context B and the brain was quickly removed. Coronal slices (300 μm thick) were cut with a vibratome (Leica VT1200S, Leica Microsystems) in artificial cerebrospinal fluid (aCSF) rich in sucrose (aCSF sucrose) containing (in mM): 2 KCl, 1.25 NaH_2_PO_4_-H_2_O, 7 MgSO_4_, 26 NaHCO_3_, 0.5 CaCl_2_, 10 glucose and 219 sucrose) at 4°C, saturated with a 95% O_2_, 5% CO_2_ mixture and maintained at pH 7.32-7.4. Then, slices were transferred to recovery chamber with a heated (35°C) oxygenated aCSF that contained (in mM) 124 NaCl, 2.5 KCl, 1.25 NaH_2_PO_4_-H_2_O, 1 MgSO_4_, 26 NaHCO_3_, 2 CaCl_2_ and 10 glucose for 15 min and incubated for > 1 h at room temperature (22 ± 2°C) and pH 7.32-7.4. Slices were transferred individually into an immersion-recording chamber and oxygenated aCSF was perfused at a rate of 2 mL/min (22 ± 2°C).

Whole-cell intracellular recordings in current clamp (CC) mode were performed in granule cell neurons in DG granule cell layer (upper horn). Cells were visualized with a water-immersion 40x objective. Patch electrodes were made from borosilicate glass capillaries (Sutter P-1000, Sutter Instruments) with resistance ranging from 4 to 6 MΩ when filled with the internal solution that contained (in mM): 130 K-MeSO_4_, 10 HEPES, 0.5 EGTA, 2 MgCl_2_, 4 Mg-ATP, 0.4 Na-GTP, 10 phosphocreatine disodium salt hydrate and 0.3 % biocytin, for membrane properties experiments in CC. KOH were used to adjust the pH of all pipette solutions to 7.2-7.3. Membrane currents and voltages were measured using Multiclamp 700B amplifiers, digitized using Digidata 1550B, and controlled using pClamp 10.7 (Molecular Devices Corporation, California, USA). Membrane intrinsic properties of granule cells were determined by applying hyperpolarizing and depolarizing current steps (1 s, with 10 pA increments from −100 to 140 pA) in CC.

## Statistical analysis

When two conditions were compared, the Shapiro-Wilks test was conducted to check the normality of the data and Fisher’s F test was used to assess the homogeneity of variances between groups. When data met the assumptions of parametric distribution, results were analyzed by unpaired student’s *t*-test. Paired *t*-tests were employed to compare paired variables. Mann-Whitney-Wilcoxon test was applied in cases where the data did not meet the requirements of normal distribution. Statistical analyses were two-tailed. For comparison between more than two groups, two-way ANOVA with different levels was conducted followed by Tukey HSD multiple comparison test. Bartlett test was used to assess the homogeneity of variances between groups. If the data distribution was non-parametric, the Kruskal-Wallis test was used followed by Mann Whitney-Wilcoxon test. All statistical analyses were two-tailed. The statistical test used is indicated in every Figure. Differences in means were considered statistically significant at p < 0.05.

Data analysis and statistics were performed using R studio (Version 1.1.463)

## Acknowledgements

Figures 1A, 2AB, 5C, E and Supplementary Figure 1A,B,E,F,J were created with BioRender.com.

---

**Supplementary Figure 1.**
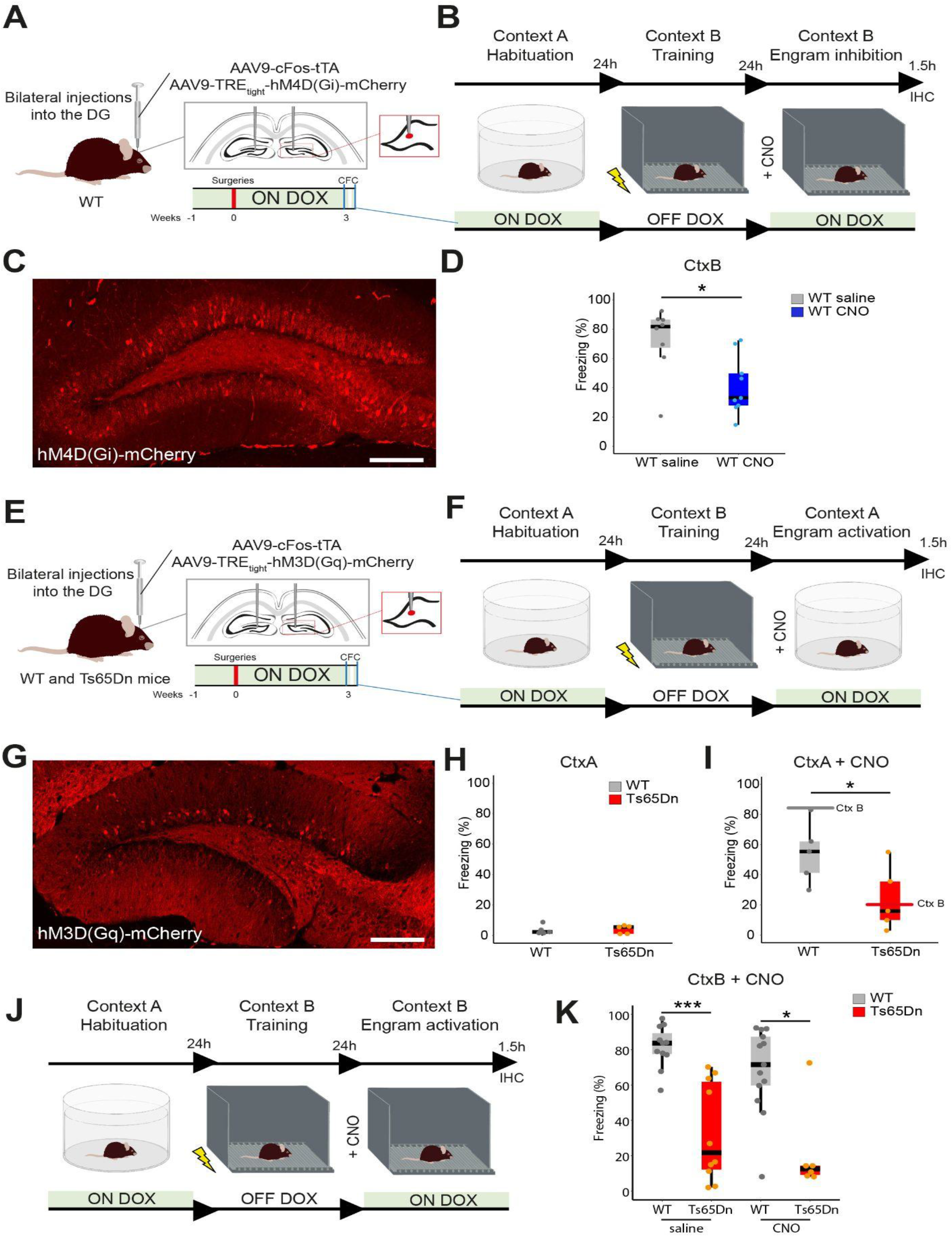
**(A-B)** Behavioral schedule used to test the inhibitory DREADD in WT. hM4D(Gi)-mCherry was expressed during training in Context B as previously described. CNO was administered 30 min before memory recall. **(C)** Representative image showing a DG section 24 h after engram-tagging procedure with hM4D(Gi)-mCherry. Scale bar = 100 μm **(D)** Engram inactivation in WT mice by CNO led to a memory impairment as evidenced by lower freezing levels when compared to saline injected controls (WT saline = 8, WT CNO = 9). Two-tailed T-test. **(E)** AAV_9_-cFos-tTA and AAV_9_-TREtight-hM3D(Gq)-mCherry viruses were injected into the dorsal DG of WT and Ts65Dn mice. **(F)** Mice were habituated in the neutral context (Context A) while ON DOX. After habituation, DOX was removed and mice were trained in Context B 24 h after. Then, the DOX diet was resumed and mice were tested again in the trained Context B after 24 h. Mice were sacrificed after 1.5 h and brains were extracted and processed for IEG quantification. **(G)** Representative image showing a DG section, 24 h after engram-tagging with hM3D(Gq)-mCherry. Scale bar = 100 μm. **(H)** Percentage of freezing of WT and Ts65Dn in Context A (WT= 5, Ts65Dn = 5). Two-tailed T-test. **(I)** Percentage of freezing of WT and Ts65Dn upon CNO (1 mg/kg) administration 30 min before the exposure to neutral Context A. Ts65Dn mice froze significantly less than WT mice (WT= 5, Ts65Dn = 5; Two-tailed T-test) Horizontal lines indicate the freezing values of WT and Ts65Dn, respectively, in CtxB (trained context). **(J)** Behavioral schedule used for testing engram activation. Engram was tagged in during the training session in Context B. CNO was administered 30 min before reexposing the mice to the trained context (B) 24 h after the training. **(K)** Percentage of freezing of WT and Ts65Dn (saline-injected) and WT and Ts65Dn mice (CNO-injected). Memory deficits in Ts65Dn were not rescued by CNO as evidenced by the reduced freezing levels compared to WT mice (WT saline = 11, Ts65Dn saline = 10, WT CNO = 13, Ts65Dn = 6). Horizontal lines indicate the freezing values of WT and Ts65Dn, respectively, in CtxB injected with saline. On the boxplots, the horizontal line indicates the median, the box indicates the first to third quartile of expression and whiskers indicate 1.5 × the interquartile range. \**P* < 0.05, ** *P* < 0.01.

**Supplementary Figure 2.**
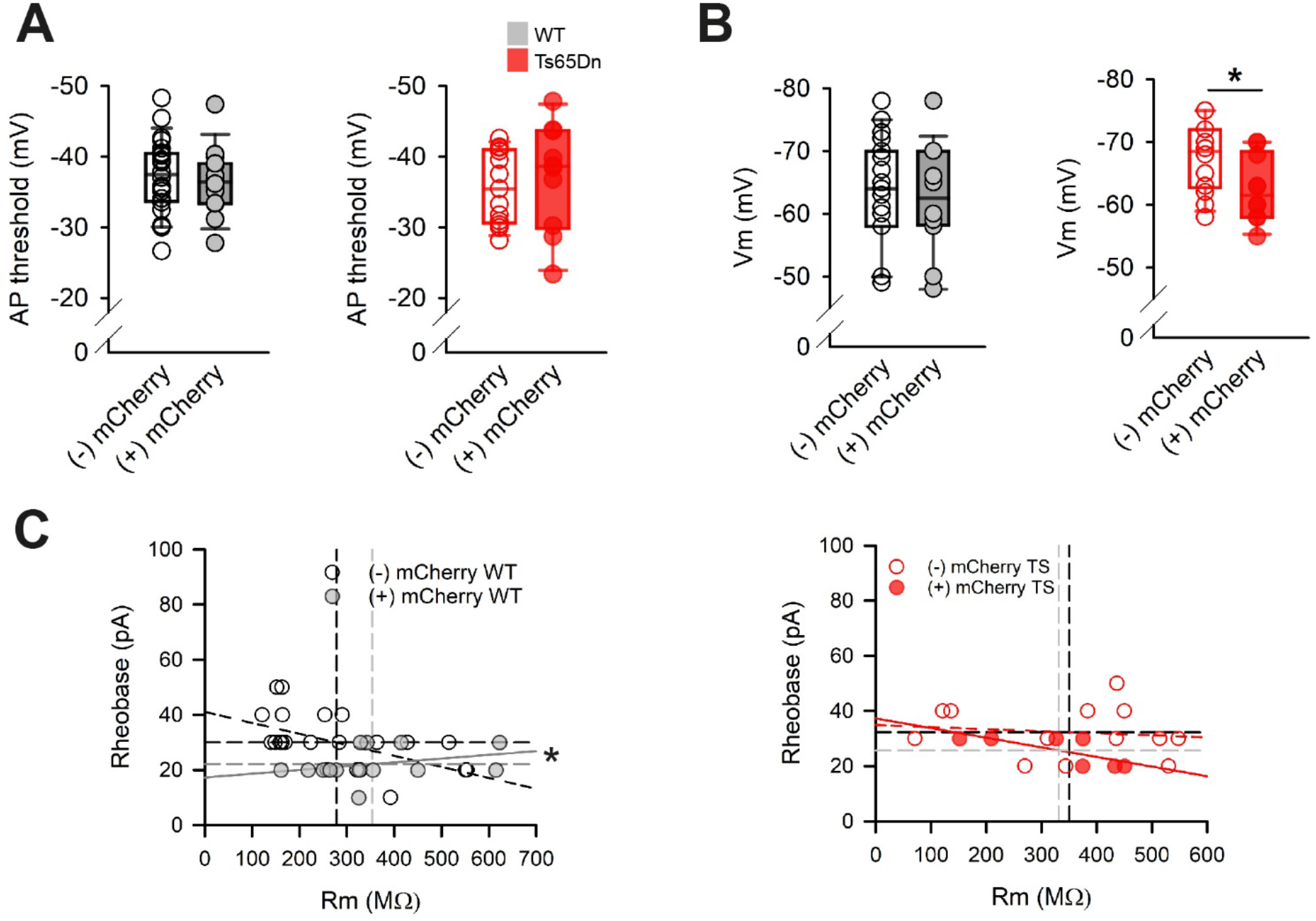
**(A)** Left: AP threshold of non-engram (black non-filled dots) and engram (gray filled dots) in WT mice (WT non-engram = 25 cells from 8 mice, WT engram = 16 cells from 8 mice). Right: AP threshold of non-engram (red non-filled dots) and engram (red filled dots) in Ts65Dn mice (Ts65Dn non-engram = 14 cells from 6 mice, Ts65Dn engram = 10 cells from 6 mice). **(B)** Left: resting membrane potential of nonengram (black non-filled dots) and engram (gray filled dots) in WT mice. (WT non-engram = 25 cells from 8 mice, WT engram = 16 cells from 8 mice). Right: resting membrane potential of non-engram (red non-filled dots) and engram (red filled dots) in Ts65Dn mice. (Ts65Dn non-engram = 14 cells from 6 mice, Ts65Dn engram = 10 cells from 6 mice). Mann Whitney. \**P* >0.05. **(C)** Left: rheobase vs. membrane resistance of non-engram (black non-filled dots) vs. engram (dark gray filled dots) in WT mice. (WT non-engram = 25 cells from 8 mice, WT engram = 16 cells from 6 mice). Right: rheobase vs. membrane resistance of non-engram (red non-filled dots) vs. engram (red filled dots) in Ts65Dn mice. (Ts65Dn non-engram = 14 cells from 8 mice, Ts65Dn engram = 10 cells from 5 mice). In every plot, horizontal and vertical black lines indicate the mean rheobase and Rm values, respectively of non-engram cells. Gray lines indicate the mean rheobase and Rm values of engram cells. Mann Whitney. \**P* >0.05. On the boxplots, the horizontal line indicates the median, the box indicates the first to third quartile of expression and whiskers indicate 1.5 × the interquartile range.

**Supplementary Figure 3.**
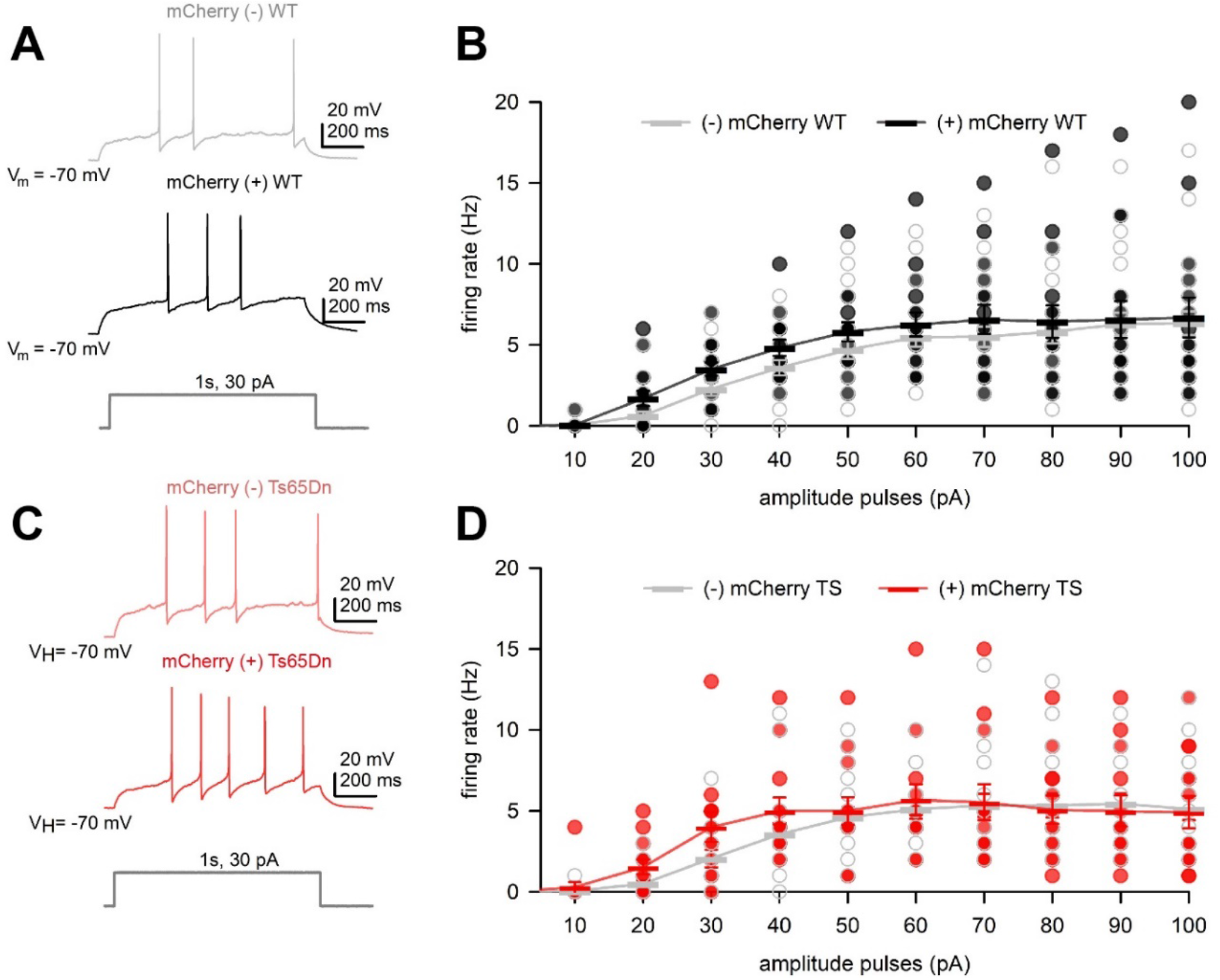
**(A)** Representative traces showing the voltage response of non-engram (gray) and engram (black) to a depolarizing current pulse in WT mice (WT non-engram = 26 cells from 8 mice, WT engram = 16 cells from 8 mice). **(B)** Current injection vs. firing rate recorded in non-engram (gray non-filled dots) and engram cells (black filled dots) in WT mice. **(C)** Representative traces showing the voltage response of non-engram (pink) and engram (red) to a depolarizing current pulse in Ts65Dn mice. (Ts65Dn non-engram = 18 cells from 7 mice, Ts65Dn engram = 13 cells from 6 mice). **(D)** Current injection vs. firing rate recorded in non-engram (gray non-filled dots) and engram cells (red filled dots) in Ts65Dn mice. ANOVA repeated measures. Data expressed as mean ± SEM.

## References

1. de la Torre R, de Sola S, Hernandez G, Farré M, Pujol J, Rodriguez J, et al. Safety and efficacy of cognitive training plus epigallocatechin-3-gallate in young adults with Down’s syndrome (TESDAD): A double-blind, randomised, placebo-controlled, phase 2 trial. Lancet Neurol. 2016;

2. Lott IT, Dierssen M. Cognitive deficits and associated neurological complications in individuals with Down’s syndrome. Vol. 9, The Lancet Neurology. 2010. p. 623–33.

3. Dierssen M. Down syndrome: the brain in trisomic mode. Nat Rev Neurosci. 2012 Dec 20;13(12):844–58.

4. Reijmers LG, Perkins BL, Matsuo N, Mayford M. Localization of a stable neural correlate of associative memory. Science (1979). 2007;

5. Liu X, Ramirez S, Pang PT, Puryear CB, Govindarajan A, Deisseroth K, et al. Optogenetic stimulation of a hippocampal engram activates fear memory recall. Nature. 2012;

6. Han JH, Kushner SA, Yiu AP, Cole CJ, Matynia A, Brown RA, et al. Neuronal competition and selection during memory formation. Science (1979). 2007;

7. Ryan TJ, Roy DS, Pignatelli M, Arons A, Tonegawa S. Engram cells retain memory under retrograde amnesia. Science (1979). 2015;

8. Pignatelli M, Ryan TJ, Roy DS, Lovett C, Smith LM, Muralidhar S, et al. Engram Cell Excitability State Determines the Efficacy of Memory Retrieval. Neuron. 2019;

9. Roy DS, Arons A, Mitchell TI, Pignatelli M, Ryan TJ, Tonegawa S. Memory retrieval by activating engram cells in mouse models of early Alzheimer’s disease. Nature. 2016 Mar 24;531(7595):508–12.

10. Hartley T, Bird CM, Chan D, Cipolotti L, Husain M, Varga-Khadem F, et al. The hippocampus is required for short-term topographical memory in humans. Hippocampus. 2007;17(1):34–48.

11. Winocur G, Moscovitch M. Memory transformation and systems consolidation. J Int Neuropsychol Soc. 2011;17(5):766–80.

12. Debiec J, LeDoux JE, Nader K. Cellular and systems reconsolidation in the hippocampus. Neuron. 2002 Oct 24;36(3):527–38.

13. Li J, Jiang RY, Arendt KL, Hsu YT, Zhai SR, Chen L. Defective memory engram reactivation underlies impaired fear memory recall in Fragile X syndrome. Elife. 2020 Oct 1;9:1–20.

14. Yiu AP, Rashid AJ, Josselyn SA. Increasing CREB function in the CA1 region of dorsal hippocampus rescues the spatial memory deficits in a mouse model of Alzheimer’s disease. Neuropsychopharmacology. 2011 Oct;36(11):2169–86.

15. Costa ACS, Scott-McKean JJ, Stasko MR. Acute injections of the NMDA receptor antagonist memantine rescue performance deficits of the Ts65Dn mouse model of Down syndrome on a fear conditioning test. Neuropsychopharmacology. 2008 Jun;33(7):1624–32.

16. Kleschevnicov AM, Belichenko P V., Villar AJ, Epstein CJ, Malenka RC, Mobley WC. Hippocampal long-term potentiation suppressed by increased inhibition in the Ts65Dn mouse, a genetic model of Down syndrome. J Neurosci. 2004 Sep 15;24(37):8153–60.

17. Siarey RJ, Carlson EJ, Epstein CJ, Balbo A, Rapoport SI, Galdzicki Z. Increased synaptic depression in the Ts65Dn mouse, a model for mental retardation in Down syndrome. Neuropharmacology. 1999;38(12):1917–20.

18. Uguagliati B, Al-Absi AR, Stagni F, Emili M, Giacomini A, Guidi S, et al. Early appearance of developmental alterations in the dendritic tree of the hippocampal granule cells in the Ts65Dn model of Down syndrome. Hippocampus. 2021 Apr 1;31(4):435–47.

19. Emili M, Stagni F, Salvalai ME, Uguagliati B, Giacomini A, Albach C, et al. Neonatal therapy with clenbuterol and salmeterol restores spinogenesis and dendritic complexity in the dentate gyrus of the Ts65Dn model of Down syndrome. Neurobiol Dis. 2020 Jul 1;140.

20. Zorrilla de San Martin J, Delabar JM, Bacci A, Potier MC. GABAergic over-inhibition, a promising hypothesis for cognitive deficits in Down syndrome. Free Radic Biol Med. 2018 Jan 1;114:33–9.

21. Fernandez F, Garner CC. Over-inhibition: a model for developmental intellectual disability. Trends Neurosci. 2007 Oct;30(10):497–503.

22. Altafaj X, Martín ED, Ortiz-Abalia J, Valderrama A, Lao-Peregrín C, Dierssen M, et al. Normalization of Dyrk1A expression by AAV2/1-shDyrk1A attenuates hippocampal-dependent defects in the Ts65Dn mouse model of Down syndrome. Neurobiol Dis. 2013 Apr;52:117–27.

23. Kleschevnikov AM, Yu J, Kim J, Lysenko L V., Zeng Z, Yu YE, et al. Evidence that increased Kcnj6 gene dose is necessary for deficits in behavior and dentate gyrus synaptic plasticity in the Ts65Dn mouse model of Down syndrome. Neurobiol Dis. 2017;

24. Smith-Hicks CL, Cai P, Savonenko A V., Reeves RH, Worley PF.. Increased sparsity of hippocampal CA1 neuronal ensembles in a mouse model of down syndrome assayed by Arc expression. Front Neural Circuits. 2017;

25. Sierra C, SM, FBA, & DM. The lncRNA Snhg11 is required for synaptic function, neurogenesis and memory and is downregulated in the dentate gyrus of Down syndrome mouse models. Res Sq. 2022;

26. Phillips RG, LeDoux JE. Differential contribution of amygdala and hippocampus to cued and contextual fear conditioning. Behavioral neuroscience. 1992;106(2):274–85.

27. Lever C, Wills T, Cacucci F, Burgess N, O’Keefe J. Long-term plasticity in hippocampal place-cell representation of environmental geometry. Nature. 2002 Mar 7;416(6876):90–4.

28. Nakashiba T, Cushman JD, Pelkey KA, Renaudineau S, Buhl DL, McHugh TJ, et al. Young dentate granule cells mediate pattern separation, whereas old granule cells facilitate pattern completion. Cell. 2012 Mar 30;149(1):188–201.

29. Pinter JD, Eliez S, Schmitt JE, Capone GT, Reiss AL. Neuroanatomy of Down’s syndrome: A high-resolution MRI study. American Journal of Psychiatry. 2001;158(10):1659–65.

30. Raz N, Torres IJ, Briggs SD, Spencer WD, Thornton AE, Loken WJ, et al. Selective neuroanatomic abnormalities in down’s syndrome and their cognitive correlates: Evidence from mri morphometry. Neurology. 1995;45(2):356–66.

31. Insausti AM, Megías M, Crespo D, Cruz-Orive LM, Dierssen M, Vallina TF, et al. Hippocampal volume and neuronal number in Ts65Dn mice: A murine model of Down syndrome. Neurosci Lett. 1998 Sep 3;253(3):175–8.

32. Navarro-Romero A, Vázquez-Oliver A, Gomis-González M, Garzón-Montesinos C, Falcón-Moya R, Pastor A, et al. Cannabinoid type-1 receptor blockade restores neurological phenotypes in two models for Down syndrome. Neurobiol Dis. 2019 May 1;125:92–106.

33. Bianchi P, Ciani E, Guidi S, Trazzi S, Felice D, Grossi G, et al. Early pharmacotherapy restores neurogenesis and cognitive performance in the Ts65Dn mouse model for down syndrome. Journal of Neuroscience. 2010 Jun 30;30(26):8769–79.

34. Leutgeb JK, Leutgeb S, Moser MB, Moser EI. Pattern separation in the dentate gyrus and CA3 of the hippocampus. Science. 2007 Feb 16;315(5814):961–6.

35. Bernier BE, Lacagnina AF, Ayoub A, Shue F, Zemelman B V., Krasne FB, et al. Dentate Gyrus Contributes to Retrieval as well as Encoding: Evidence from Context Fear Conditioning, Recall, and Extinction. J Neurosci. 2017;37(26):6359–71.

36. Poll S, Mittag M, Musacchio F, Justus LC, Giovannetti EA, Steffen J, et al. Memory trace interference impairs recall in a mouse model of Alzheimer’s disease. Nat Neurosci. 2020 Aug 1;23(8):952–8.

37. Zhang Z, Ferretti V, Güntan I, Moro A, Steinberg EA, Ye Z, et al. Neuronal ensembles sufficient for recovery sleep and the sedative actions of α2 adrenergic agonists. Nat Neurosci. 2015 Apr 28;18(4):553–61.

38. Tonegawa S, Liu X, Ramirez S, Redondo R. Memory Engram Cells Have Come of Age. Neuron. 2015 Sep 2;87(5):918–31.

39. Cramer NP, Xu X, Haydar TF, Galdzicki Z. Altered intrinsic and network properties of neocortical neurons in the Ts65Dn mouse model of Down syndrome. Physiol Rep. 2015;3(12).

40. Stern S, Segal M, Moses E. Involvement of Potassium and Cation Channels in Hippocampal Abnormalities of Embryonic Ts65Dn and Tc1 Trisomic Mice. EBioMedicine. 2015 Sep 1;2(9):1048–62.

41. Dierssen M, Benavides-Piccione R, Martínez-Cué C, Estivill X, Flórez J, Elston GN, et al. Alterations of neocortical pyramidal cell phenotype in the Ts65Dn mouse model of Down syndrome: Effects of environmental enrichment. Cerebral Cortex. 2003;

42. Yiu AP, Rashid AJ, Josselyn SA. Increasing CREB Function in the CA1 Region of Dorsal Hippocampus Rescues the Spatial Memory Deficits in a Mouse Model of Alzheimer’s Disease. Neuropsychopharmacology 2011 36:11. 2011 Jul 6;36(11):2169–86.

43. Hyde LA, Frisone DF, Crnic LS. Ts65Dn mice, a model for Down syndrome, have deficits in context discrimination learning suggesting impaired hippocampal function. Behavioural Brain Research. 2001 Jan 8;118(1):53–60.

44. Kleschevnikov AM, Belichenko P V., Faizi M, Jacobs LF, Htun K, Shamloo M, et al. Deficits in cognition and synaptic plasticity in a mouse model of Down syndrome ameliorated by GABAB receptor antagonists. J Neurosci. 2012 Jul 4;32(27):9217–27.

45. Best TK, Cramer NP, Chakrabarti L, Haydar TF, Galdzicki Z. Dysfunctional hippocampal inhibition in the Ts65Dn mouse model of Down syndrome. Exp Neurol. 2012;

46. Rao-Ruiz P, Yu J, Kushner SA, Josselyn SA. Neuronal competition: microcircuit mechanisms define the sparsity of the engram. Current Opinion in Neurobiology. 2019.

47. Miry O, Li J, Chen L. The Quest for the Hippocampal Memory Engram: From Theories to Experimental Evidence. Front Behav Neurosci. 2021 Jan 15;14.

48. Leake J, Zinn R, Corbit LH, Fanselow MS, Vissel B. Engram Size Varies with Learning and Reflects Memory Content and Precision. J Neurosci. 2021 May 5;41(18):4120–30.

49. Morrison DJ, Rashid AJ, Yiu AP, Yan C, Frankland PW, Josselyn SA. Parvalbumin interneurons constrain the size of the lateral amygdala engram. Neurobiol Learn Mem. 2016 Nov 1;135:91–9.

50. Aimone JB, Deng W, Gage FH. Resolving new memories: a critical look at the dentate gyrus, adult neurogenesis, and pattern separation. Neuron. 2011 May 26;70(4):589–96.

51. Baker S, Vieweg P, Gao F, Gilboa A, Wolbers T, Black SE, et al. The Human Dentate Gyrus Plays a Necessary Role in Discriminating New Memories. Curr Biol. 2016 Oct 10;26(19):2629–34.

52. Smith GK, Kesner RP, Korenberg JR. Dentate Gyrus Mediates Cognitive Function in the Ts65Dn/DnJ Mouse Model of Down Syndrome. Hippocampus. 2014 Mar 1;24(3):354.

53. Josselyn SA, Tonegawa S. Memory engrams: Recalling the past and imagining the future. Science. 2020 Jan 3;367(6473).

54. Dong Y, Green T, Saal D, Marie H, Neve R, Nestler EJ, et al. CREB modulates excitability of nucleus accumbens neurons. Nat Neurosci. 2006;

55. Zhou Y, Won J, Karlsson MG, Zhou M, Rogerson T, Balaji J, et al. CREB regulates excitability and the allocation of memory to subsets of neurons in the amygdala. Nat Neurosci. 2009 Nov;12(11):1438–43.

56. Sekeres MJ, Neve RL, Frankland PW, Josselyn SA. Dorsal hippocampal CREB is both necessary and sufficient for spatial memory. Learn Mem. 2010 Jun;17(6):280–3.

57. Popov VI, Kleschevnikov AM, Klimenko OA, Stewart MG, Belichenko P V. Three-dimensional synaptic ultrastructure in the dentate gyrus and hippocampal area CA3 in the Ts65Dn mouse model of Down syndrome. J Comp Neurol. 2011 May 1;519(7):1338–54.

58. Guidi S, Stagni F, Bianchi P, Ciani E, Ragazzi E, Trazzi S, et al. Early pharmacotherapy with fluoxetine rescues dendritic pathology in the Ts65Dn mouse model of down syndrome. Brain Pathol. 2013 Mar;23(2):129–43.

